# Natural competence promotes high rates of DNA transfer between strains of the core bee gut bacterium *Snodgrassella alvi*

**DOI:** 10.64898/2025.12.12.693966

**Authors:** A H M Zuberi Ashraf, Kadena K. Weaver, Patrick J. Lariviere, Jeffrey E. Barrick

**Author notes:** Systems, Synthetic, and Physical Biology Ph.D. Program, Rice University, Houston, Texas, USA (K.K.W.); Department of Biology, University of Miami, Coral Gables, Florida, USA (P.J.L.); Departments of Microbiology, Genetics, & Immunology and Entomology, Michigan State University, East Lansing, Michigan, USA (J.E.B.).

## Abstract

Sexual recombination and horizontal gene transfer are expected to improve the survival of host-associated bacteria that face colonization bottlenecks and intense competition. We serendipitously observed recombination between *Snodgrassella alvi* strains within bees, which led us to discover that this core member of the bee gut microbiota is naturally competent, including under laboratory conditions. High rates of gene transfer via DNA release and uptake by *S. alvi* have implications for how it has evolved in managed hives and suggest opportunities for using *in situ* microbiome engineering to protect bee health.

---

Honey bees (*Apis mellifera*) have a co-evolved hindgut microbial community with several core bacterial species^1,2^. These bacteria are acquired by worker bees through fecal-oral transmission and contact with hive surfaces. *Snodgrassella alvi* is a primary colonizer that forms a biofilm directly on the epithelium of the ileum^3,4^. It stimulates the host immune system and has metabolic interactions with the host and other core members of the microbiota, including *Gilliamella apis*^*5*^. Multiple *S. alvi* strains circulate in a hive, and more than one may colonize the same bee^6,7^. *S. alvi* possess type VI secretion systems and a diverse repertoire of effectors^8^. Some of these have been shown to mediate competition between different *S. alvi* strains within the bee gut^9^.

To observe how this bee gut symbiont evolves in its host environment, we passaged three populations of *S. alvi* wkB2 through four cohorts of microbiota-deficient adult bees over 28 days (**Fig. 1a**). Each population was initiated by feeding an *S. alvi* culture grown from a different bacterial colony to a group of newly emerged bees that was separately caged in the laboratory. Thereafter, each *S. alvi* population was transferred to the next cohort of newly emerged adult bees every seven days by feeding gut homogenate from the previous cohort. In two of the three *S. alvi* populations, we found clusters of mutations dominated by synonymous base substitutions in the genomes of clones isolated at the conclusion of the experiment. These mutations were spread across either a 5.4-kbp region—which includes σ^54^ -dependent regulator (*RS08390*), HAMP domain-containing histidine kinase (*RS08395*), and topoisomerase IV subunit A (*parC*) genes— or a larger 22-kbp region that overlaps the smaller region and includes additional genes (**Fig. 1b**). This mutational signature is a hallmark of sexual recombination transferring alleles from a related *S. alvi* strain.

**Fig. 1.**
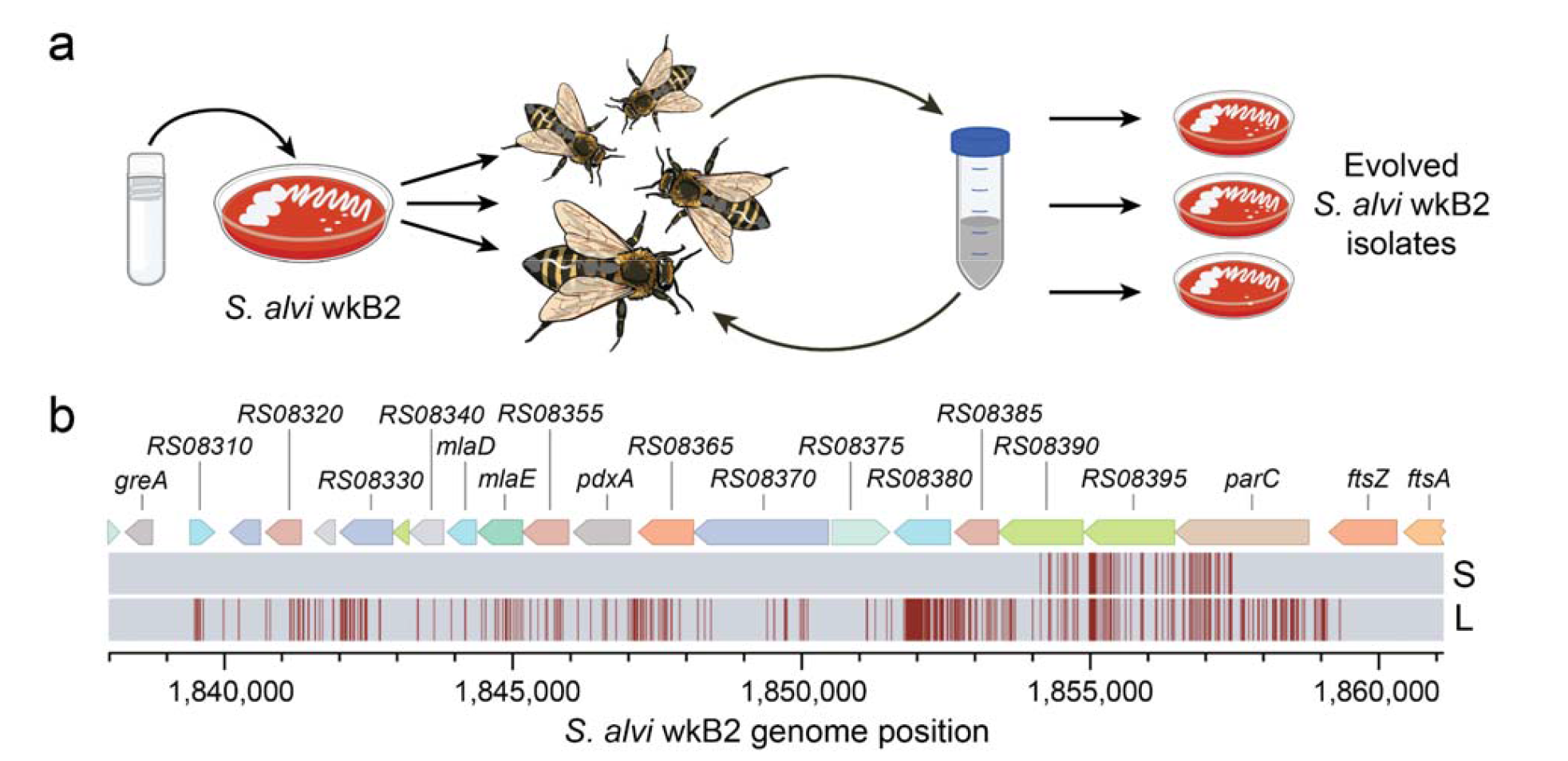
Recombination events serendipitously observed between *S. alvi* in the honey bee gut. **a**, Three separate populations of *S. alvi* wkB2 initiated from clonal isolates were fed to cohorts of microbiome-deficient bees and allowed to colonize their guts for a week. Gut homogenate from these bees was fed to new cohorts of bees. After four total colonization cycles, gut homogenate was plated to isolate evolved bacterial clones. **b**, Two different recombination events were observed in the genomes of *S. alvi* isolated from two of the three evolution experiment populations. Alleles from an unknown, hive-associated *S. alvi* strain (vertical red bars) within a small (S) or overlapping large (L) region were transferred to the focal wkB2 strain. Full lists of mutations are provided in **Supplementary Table 1**.

The guts of microbiota-deficient bees that emerge in the laboratory can contain some microbes from their hive of origin, though the titer of bacteria typically remains very low if they are not experimentally colonized^10^. While we only isolated derivatives of the original wkB2 strain we used for colonization when picking endpoint clones to analyze, the bees that we used appear to have also harbored an unknown *S. alvi* strain. That overlapping genomic regions were acquired by our focal strain from this hive-associated *S. alvi* strain in at least two independent instances suggests that these recombination events were beneficial.

Further sequencing of the ancestral wkB2 stock revealed that the two populations with recombination events were derived from a subpopulation of cells that had acquired a single-base insertion causing a frameshift in the gene *RS08395* during prior growth in the laboratory. It is possible that the recombination events were selected for because they restored function of this HAMP domain-containing histidine kinase gene and that this was beneficial in the bee gut, though a transposon insertion sequencing study found only a minor defect in fitness for loss-of-function mutations in this gene that was not statistically significant^5^. Alternatively or in addition, the recombination events may have transferred other beneficial alleles. We did not detect any other new mutations elsewhere in the genomes of the evolved clones that experienced recombination.

To understand how commonly similar recombination events occur between *S. alvi* strains within the bee gut, we co-colonized microbiota-deprived bees with two marked strains. Each strain was engineered to have a different antibiotic resistance gene inserted at a unique location in its genome. After 5 days, we plated the gut contents of these bees and counted colony forming units (CFUs) on agar with and without these antibiotics. We found that the frequencies of doubly resistant CFUs—representing cells that had acquired the antibiotic resistance gene from the other strain—relative to total CFUs were 0.9×10^−4^ and 5.7×10^−4^ in two separate experimental trials using different cohorts of bees (**Fig. 2a**), confirming that recombination occurred at a high rate *in vivo*.

**Fig. 2.**
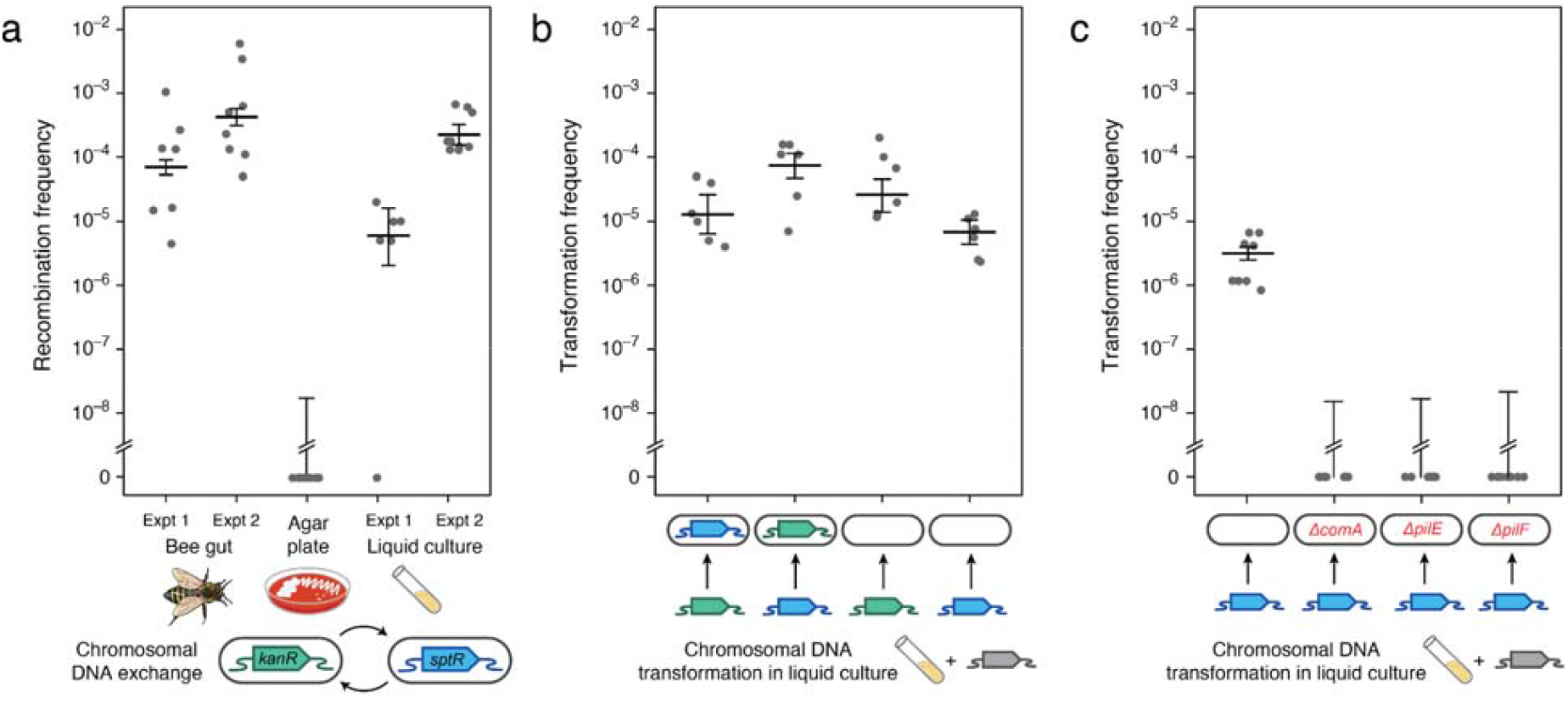
*S. alvi* wkB2 is naturally competent. **a**, Gene transfer occurs between *S. alvi* strains when they co-colonize the bee gut or are co-cultured in a liquid medium but not on an agar plate. Two *S. alvi* wkB2 strains marked by integrating genes conferring kanamycin (Kan^r^) or spectinomycin (Spt^r^) resistance into their chromosomes at different locations were mixed equally to begin each experiment. *S. alvi* wkB2 is naturally resistant to tetracycline (Tet^r^). Recombination frequency was measured as CFUs on selective Kan+Spt agar relative to CFUs on nonselective Tet agar. Two trials of the bee colonization and liquid culture experiments were conducted at different times. **b**, *S. alvi* strains can acquire genes conferring Kan^r^ or Spt^r^ phenotypes from chromosomal DNA. Purified DNA from one marked *S. alvi* strain was added to a liquid culture of the opposite marked strain or wild-type *S. alvi*. Transformation frequency was measured as CFUs on selective plates relative to CFUs on nonselective plates. **c**, Disrupting genes that encode predicted components of the competence machinery eliminates gene acquisition. Experiments were performed as in **b** to compare acquisition of Spt^r^ from DNA purified from a strain with the marker gene integrated into its chromosome. Wild-type *S. alvi* wkB2 and strains with mutations inactivating *comA, pilE*, and *pilF* were tested. In all panels, points are biological replicates representing individual bees or cultures, horizontal bars are maximum likelihood estimates of transformation frequencies, and error bars are 95% confidence intervals estimated using Poisson regression (see **Methods**).

Next, we investigated whether recombination could occur *in vitro. S. alvi* is typically grown in the laboratory on a solid agar medium. Co-culturing mixtures of the marked *S. alvi* strains on these agar plates did not yield any doubly resistant colonies (**Fig. 2a**). However, when we co-cultured these strains in a liquid medium we found recombinants at frequencies that were similar to those observed in the bee gut: 0.16×10^−4^ and 3.23×10^−4^ in two separate experimental trials (**Fig. 2a**). Whole-genome sequencing of putative recombinants confirmed transfer of antibiotic resistance genes had occurred.

Liquid cultures of a singly marked strain or wild-type wkB2 were also able to acquire the antibiotic resistance genes from chromosomal DNA isolated from a marked strain (**Fig. 2b**). We detected DNA in the supernatant of liquid *S. alvi* cultures and found that its concentration increased over time (**Extended Data Fig. 1**). These observations are consistent with extracellular DNA released from *S. alvi* cells, possibly during biofilm formation, being directly taken up by other cells through natural competence.

*S. alvi* is in the *Neisseriaciae* family, and strain wkB2 encodes genes homologous to all components of the *Neisseria gonorrhea* competence machinery (**Supplementary Table 2**). ComA is an inner membrane DNA transporter, PilE is the major pilin subunit that forms the type IV competence pilus^5^, and PilF is involved in pilus assembly^11^. Disrupting the *S. alvi* gene encoding any of these three proteins completely eliminated acquisition of an antibiotic resistance gene from marked chromosomal DNA (**Fig. 2c**), confirming that *S. alvi* is naturally competent. *Neisseria* species have copies of a species-specific DNA uptake sequence (DUS) spread throughout their genomes. *S. alvi* genomes were previously noted to not contain any such repeats that would be a candidate DUS^5^. DNA uptake by *S. alvi* appears to be sequence nonspecific, given that we have also engineered its genome by using natural transformation to integrate constructs from *E. coli* plasmids containing only 1000-bp flanking homology regions (see **Methods**).

Competence for DNA uptake is regulated by a variety of environmental cues in other microbes. Some bacteria, like *Acinetobacter baylyi*, are constitutively competent, whereas others, like *Streptococcus pneumoniae* and *Bacillus subtilis*, are only competent during specific growth phases or in certain environments^12^. For *S. alvi*, we found that over a 72-hour growth period, transformation rates were highest in the first 24 hours, when cells were actively dividing, before reaching a saturating density at around 48 hours (**Extended Data Fig. 2**). Another naturally competent species, *Vibrio cholerae*, activates DNA uptake in response to growth on chitin^13^. The gut lining of insects, including honey bees, contains chitin, but this potential environmental cue did not have a consistent effect on *S. alvi* competence in liquid cultures (**Extended Data Fig. 3**).

In previous work, we and others have used electroporation of DNA constructs to modify the *S. alvi* chromosome^4,11,14^. To determine whether DNA uptake via natural competence was responsible for the success of this approach, we compared transformation of wkB2 with and without electroporation. We tested two types of DNA. The first was a non-replicating plasmid containing an antibiotic resistance gene cassette with 1000-bp flanking homology to the *S. alvi* chromosome on each side. The second was DNA from an *S. alvi* strain with the same antibiotic resistance cassette integrated into its chromosome. Electroporation yielded 3.7-fold and 7.6-fold increases in transformation frequencies, respectively, for plasmid and chromosomal DNA (**Extended Data Fig. 4**). Thus, while electroporation does improve *S. alvi* transformation to some extent, natural transformation should be sufficient for most routine genetic engineering procedures.

Prior comparative genomics studies found evidence of widespread recombination between *S. alvi* honey bee strains^7,15^, but the mechanism was unknown. Here, we experimentally confirmed that *S. alvi* strains readily exchange DNA and integrate extracellular DNA. The natural competence of *S. alvi* has implications for its evolution and ecology as a host-restricted symbiont. It enables sexual recombination, which can preserve and improve fitness, particularly in the face of colonization bottlenecks which might otherwise lead to the accumulation of deleterious mutations. Natural competence can also catalyze the acquisition of genes that mediate antagonistic interactions within microbiomes. *V. cholerae*^*16*^ and *A. baylyi*^*17*^ use type VI secretion system effectors to lyse competitors and can then take up their DNA. In the bee gut, the ability to acquire new effectors and immunity proteins through natural transformation would be expected to give one *S. alvi* strain a competitive advantage over others.

Oxytetracycline is commonly used in beekeeping to treat foulbrood disease due to its broad-spectrum effects against bacteria and low toxicity to bees^18^. Many honey bee *S. alvi* strains have become resistant to oxytetracycline by acquiring genes that encode efflux pumps or block antibiotic binding to the ribosome^19,20^. It is possible that natural transformation facilitated the spread of oxytetracycline resistance among *S. alvi* strains once it was acquired by some. This could have preserved more overall genetic diversity within these bacteria in managed bee colonies than there would have been if there had been strong sweeps of a few initially resistant asexual lineages of *S. alvi* that acquired resistance through other gene transfer mechanisms, such as conjugation^19^.

High rates of natural transformation have implications for proposed uses of *S. alvi* in microbiome engineering applications^21^, such as using symbiont-mediated RNAi to protect bees from parasites and pathogens^22^. On one hand, natural competence complicates biocontainment. Even within one bee, native *S. alvi* strains would be expected to frequently acquire DNA sequences from an engineered strain. On the other hand, this capability might be used to purposefully transfer a DNA construct into native *S. alvi* strains from an engineered donor. This approach could disseminate beneficial traits into strains that already exist in a given bee colony rather than requiring that a foreign strain displace them within the hive microbiome. Persistence of the engineered function could be controlled by linking it to genes for utilizing an orthogonal nutrient or other selectable phenotype. It may even be possible to feed DNA or DNA-containing nanoparticles or vesicles to bees to directly engineer their native *S. alvi*. Future work can establish the feasibility and safety of these *in situ* engineering approaches.

## Methods

### Culture conditions

*Snodgrassella alvi* wkB2 was cultured on Columbia agar (BD Difco) with 5% sheep blood (VWR) (CBA) or in Columbia broth (BD Difco). Unless otherwise specified, *S. alvi* cultures and plates were incubated for 48 h at 35°C with 5% CO_2_. For growth in liquid media, cultures were shaken at 200 r.p.m. Media were supplemented with spectinomycin (Spt) at 30 µg/ml and kanamycin (Kan) at 25 μg/ml where specified.

Tetracycline (Tet) was added to some media at 10 μg/ml to prevent contamination, as *S. alvi* wkB2 is naturally resistant to this antibiotic. Where indicated, cells were washed and resuspended in phosphate-buffered saline (PBS) (Fisher). Strains and plasmids used in this study are described in **Supplementary Table 3**.

### Collecting and rearing honey bees

Bees were from hives maintained by the laboratory of Nancy Moran on the University of Texas at Austin campus. Microbiota-deficient newly emerged worker bees were obtained by allowing pupae to develop and emerge from frames in the laboratory by chewing out through the caps. Microbiota-deprived bees were obtained by pulling pupae from frames using sterilized tweezers and placing them in a vented box to complete development. Adult bees were reared in cup cages. In all cases, bees were kept in environmental chambers at 34.5°C and 80% relative humidity in the dark.

### Colonizing honey bees

Sugar syrup was prepared by mixing sucrose in a 1:1 (w/w) ratio with deionized H_2_O and filter sterilized. An inoculum mix was prepared by resuspending *S. alvi* scraped from CBA plates in PBS at an optical density at 600 nm (OD_600_) of 0.5 and mixing it with sugar syrup in a 1:5 ratio. Groups of microbiota-deficient bees were placed in separate 50 ml conical tubes, coated with 350 μl of the bacterial suspension in sugar syrup, and allowed to allogroom for 30 minutes. Cup cages were provisioned with sterilized pollen that was also coated with 350 μl of the bacterial suspension in sugar syrup. A 10 ml conical feeding tube with holes poked in the bottom to drip sugar syrup was inserted from the top of the cup cage. Bees were transferred from the conical tubes to their respective cup cages and placed in an environmental chamber. Cup cages were checked regularly, and the sugar syrup feed was replaced as needed.

### *S. alvi* evolution experiment in bees

Individual colonies of *S. alvi* from a CBA plate were streaked out and grown on separate CBA plates to create independent initial inocula for each population of the evolution experiment. Cohorts of 10 microbiota-deficient bees were colonized with *S. alvi* from each of these agar plates as described above. After seven days, cup cages were placed at 4°C for 15 minutes to immobilize bees and three bees were picked. Their guts were dissected and homogenized with a pestle in 1 ml PBS in a microfuge tube. The pooled gut homogenate was then used to colonize a new cohort of bees over seven days. This process was repeated twice more for a total of four passages over 28 days. Portions of the gut homogenates from bees isolated after each passage were cryopreserved.

### Genome sequencing

Genomic DNA was extracted using an Invitrogen PureLink Genomic DNA Mini Kit (K182001, Invitrogen). Methods for whole-genome sequencing and analysis were as described elsewhere^11^. Briefly, mutations in endpoint clones were identified from Illumina sequencing data using *breseq* (v0.38.1)^23^.

### Genome modification

Genome modifications were performed as previously described^11^. Briefly, 1000-bp homology arms on each side of the target region were amplified using PCR. Integration vectors were assembled containing these homology arms flanking an insertion cassette containing an antibiotic resistance gene. Plasmid DNA was isolated using a QIAprep Spin Miniprep Kit (27104, QIAGEN). *S. alvi* was electroporated with one of these plasmids or with genomic DNA from a strain that had previously been engineered.

Plating on CBA+Kan or CBA+Spt was used to select for cells with the desired insertions or gene replacements in their chromosomes.

### DNA exchange and transformation assays

Honey bee co-colonization DNA exchange experiments used methods as described for the evolution experiment, except microbiota-deprived bees were used and inoculated with a 1:1 mixture of the two strains being tested. After 5 days of colonization, 5 µl spots of 10-fold serial dilutions of gut homogenate from individual bees were plated. Colony counts on CBA+Kan+Spt plates were divided by colony counts on CBA+Tet plates to determine the recombination frequencies.

For testing DNA exchange in co-culture. *S. alvi* strains were grown separately on plates, resuspended in PBS to an OD_600_ of 0.5, and combined in a 1:1 ratio to create an inoculum stock. For exchange on agar, 100 µl of inoculum stock was plated in a spot on CBA+Tet. After 72 h of incubation, cells were scraped into PBS, mixed to resuspend, then serially diluted and plated as in the co-colonization experiment. For exchange in liquid, 5 µl of inoculum stock was added to 1 ml of CB. After 72 h of incubation, the culture was serially diluted and plated in the same way.

For transformation assays in liquid cultures, an inoculum mixture was prepared by resuspending a single *S. alvi* strain grown on a CBA plate to an OD_600_ of 0.5. Then, 1 ml of CB was inoculated with 5 µl of this stock and 1 µg of chromosomal DNA was added. Chromosomal DNA was isolated from cultures of strains with no plasmids, as described above. After 72 h of incubation, each culture was serially diluted and spot plated.

Colony counts on selective CBA plates, containing antibiotics to select for resistance genes in the recipient strain (if applicable) and in the donor DNA, were divided by colony counts on nonselective CBA+Tet plates to determine transformation frequencies.

For testing the effect of chitin on DNA exchange and transformation, we adapted methods previously used with *Vibrio cholerae*^*24*^. Briefly, 80 mg of chitin powder from shrimp shells (C7170, Sigma) was measured out in a test tube and autoclaved. Then, 990 μl of CB was added. For a transformation assay, chromosomal DNA was added. Next, we inoculated with 10 μl of an *S. alvi* culture resuspended to OD_600_ 0.5 for a transformation assay or a 1:1 mixture of two strains for a DNA exchange assay. After incubation for 72 h under standard conditions, test tubes were vigorously vortexed to detach cells from the chitin followed by serial dilution and plating for colony counts.

For measuring the effect of growth phase on transformation, *S. alvi* was grown in 10 ml of CB, inoculated with 100 μl of a 0.5 OD_600_ resuspension of cells grown on CBA. At 24, 48, and 72 h, chromosomal DNA was added to 1 ml of the culture after it was transferred to a new test tube. Cultures with DNA were incubated for 2 h more, followed by adding 20 U of DNase I (M0303, New England Biolabs) and incubating for an additional 30 min. Then, samples were serially diluted and plated for colony counts.

For determining whether electroporation affected transformation, competent *S. alvi* cells were prepared and electroporated as previously described^11^. We added chromosomal DNA or plasmid DNA for a non-replicating integration vector, isolated as described above, to 75 μl competent cell aliquots. One set of samples was electroporated. Then, 900 µl of CB was added to both sets. Following incubation for 24 h, cultures were serially diluted and plated for colony counts.

Full details for each experiment—including recipient strains, strains used as DNA sources, DNA concentrations, and colony counts—are provided as **Source Data**.

### DNA exchange and transformation frequency analysis

For all experiments the number of CFUs on plates containing antibiotics selecting for recombinant cells versus the number of CFUs on nonselective plates on which the original strain or strains grew was used to estimate the frequency of exchange or transformation. When colonies were observed in at least one replicate on selective media, a Poisson generalized linear model with selective versus nonselective plate and biological replicate as fixed factors was used to estimate the maximum likelihood frequency and 95% confidence intervals. When all counts were zero, a Poisson rate model was used to estimate an upper 95% confidence limit. The statistical significance of a frequency difference between treatments was evaluated by comparing the fits of Poisson regressions with and without this additional factor using a likelihood ratio test.

### Measuring DNA release by *S. alvi*

*S. alvi* liquid cultures were started from single colonies and grown for 72 h. At each time point, 1 ml samples were removed from each culture. Cells were removed from samples through centrifugation and then passing supernatant through a 0.2 μm syringe filter. A portion of each filtered supernatant was loaded on a 1% agarose gel cast with SYBR Safe dye (S33102, Invitrogen). An aliquot of 1 kb Plus DNA Ladder (New England Biolabs) was loaded as a DNA concentration standard. After electrophoresis, gels were imaged using a Gel Doc XR+ Imager (Bio-Rad). Background-corrected band volumes were quantified using ImageJ. A standard curve was constructed from several bands of the ladder and used to estimate the amount of chromosomal DNA in each sample.

## Data availability

Whole-genome sequencing data are available from the NCBI Sequence Read Archive (Accession: PRJNA1335949). All other data supporting the findings of this study are found in the Article, Supplementary Information, and Source Data.

## Acknowledgements

We thank Kim Hammond, Eli Powell, and Nancy Moran for maintaining honey bee colonies used for this work; Kathleen Sotelo for assistance with the extracellular DNA release assay; and Isaac Gifford for suggestions regarding competence assays. This study was funded by the U.S. National Science Foundation (IOS-2103208 to J.E.B.) and the U.S. Army Research Office (W911NF-20-1-0195 to J.E.B.), and the U.S.

Department of Agriculture (2023-67012-39356 to P.J.L.). A.Z.A. acknowledges support from a University of Texas at Austin Continuing Graduate Fellowship.

## Author Contributions

Conceptualization: A.Z.A. and J.E.B.; Investigation: A.Z.A. and K.K.C.; Writing - Original Draft: A.Z.A. and J.E.B.; Writing - Review & Editing: A.Z.A., K.K.C, P.J.L., and J.E.B. Visualization: A.Z.A. and J.E.B.; Supervision: P.J.L. and J.E.B.; Funding acquisition: J.E.B.

---

**Extended Data Fig. 1.**
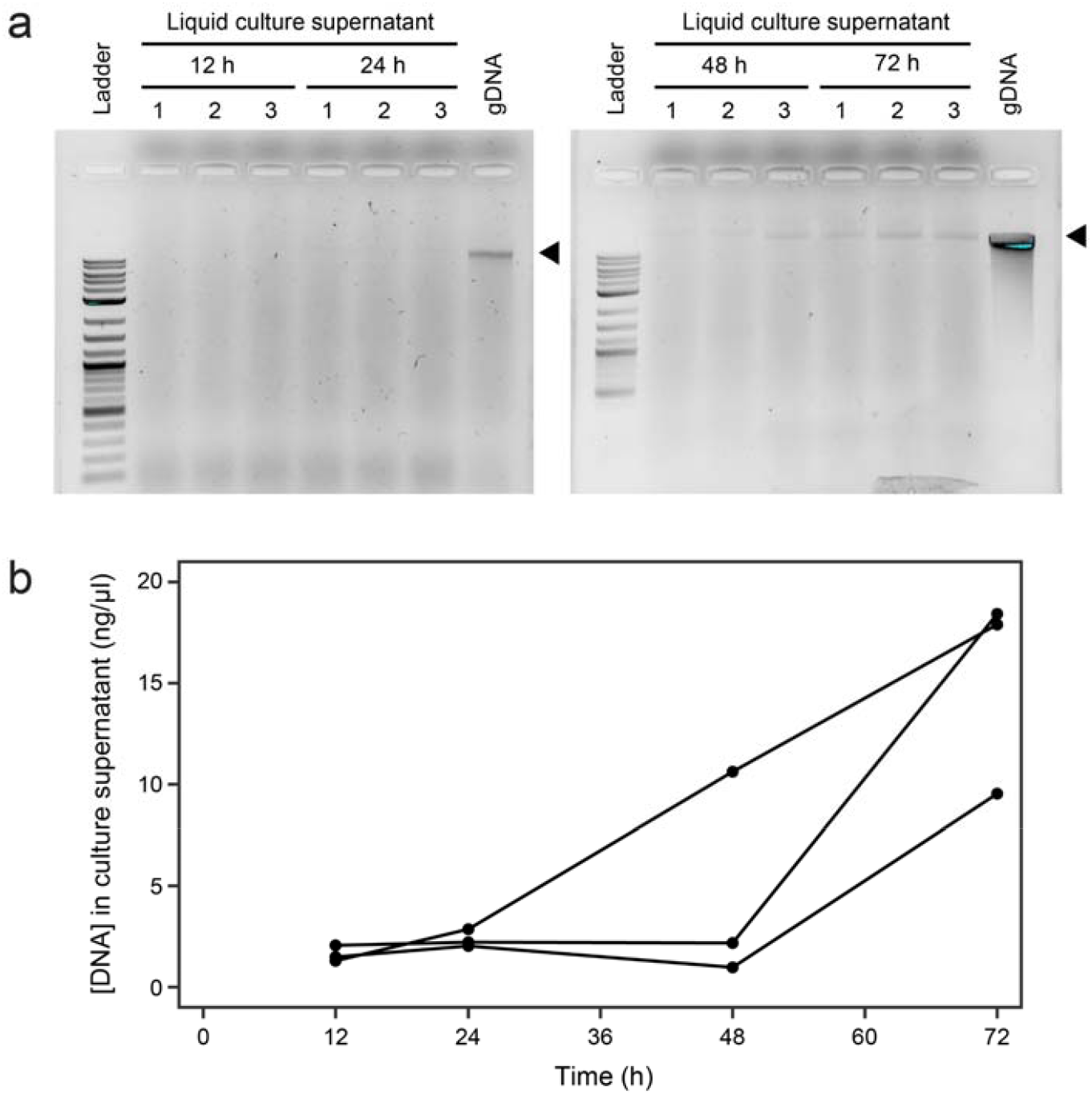
Genomic DNA accumulates in the supernatant of *S. alvi* cultures. **a**, Samples of three replicate liquid cultures of *S. alvi* (labeled 1, 2, and 3) taken over a time course (after 12, 24, 48, and 72 h) were centrifuged and filtered to remove cells before running them on an agarose gel. The band/region used to quantify chromosomal DNA is indicated with a triangle next to the lane loaded with genomic DNA isolated from *S. alvi* cells (gDNA). **b**, Estimates of chromosomal DNA amounts in each sample, determined by comparing the volume of the band/region indicated with a triangle in **a** to a standard curve constructed from several bands of the ladder. Each line tracks the change in DNA concentration over time for a different replicate culture.

**Extended Data Fig. 2.**
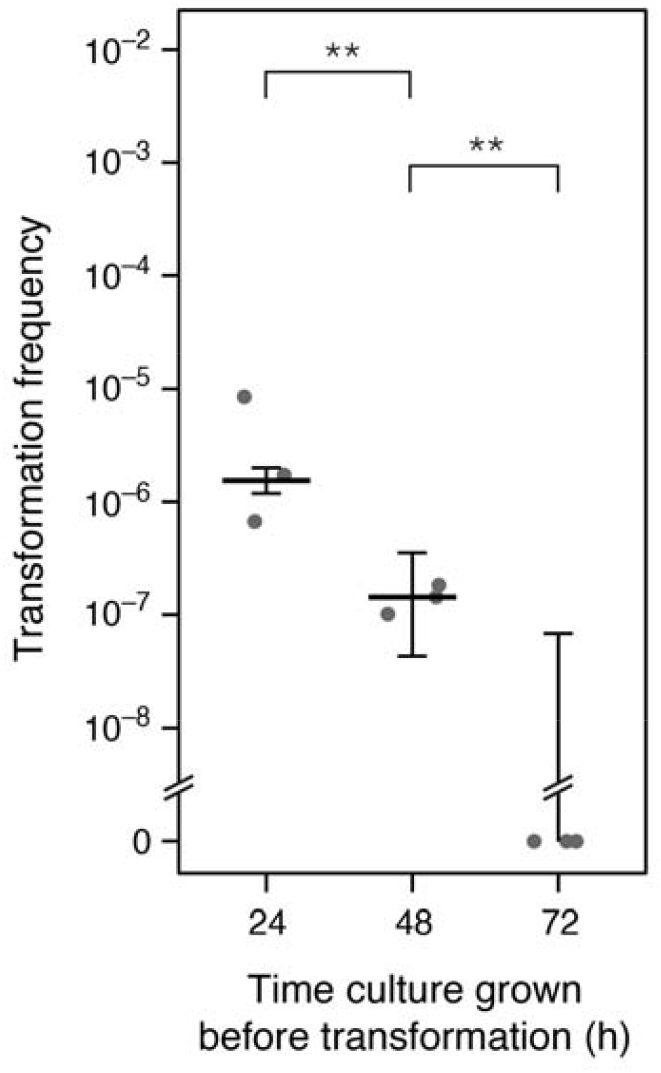
*S. alvi* transformation frequencies at different growth phases. DNA was added to aliquots of liquid cultures after growth for the specified time intervals. Transformation was allowed to proceed for 2 h before adding DNase to prevent further DNA uptake. Statistical significance for comparisons: \*\**p* < 0.01; \**p* < 0.05; ns, nonsignificant (Poisson regression likelihood ratio tests).

**Extended Data Fig. 3.**
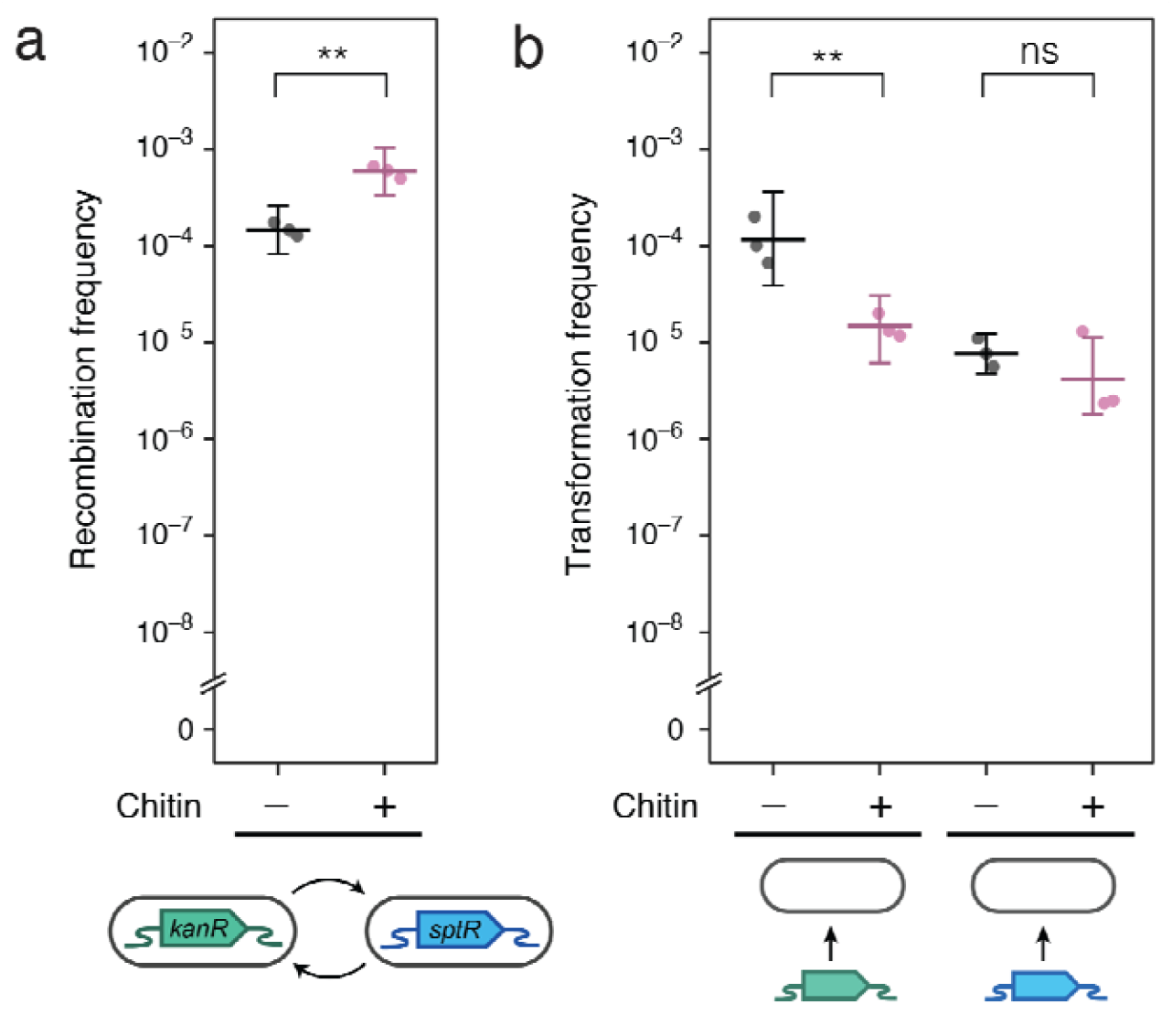
Effect of chitin on *S. alvi* transformation. **a**, DNA exchange assay testing the effect of adding chitin to liquid cultures. **b**, DNA transformation assay testing the effect of adding chitin to liquid cultures. Strains and DNA constructs were the same as in **Fig. 1**. Statistical significance for comparisons: \*\**p* < 0.01; \**p* < 0.05; ns, nonsignificant (Poisson regression likelihood ratio tests).

**Extended Data Fig. 4.**
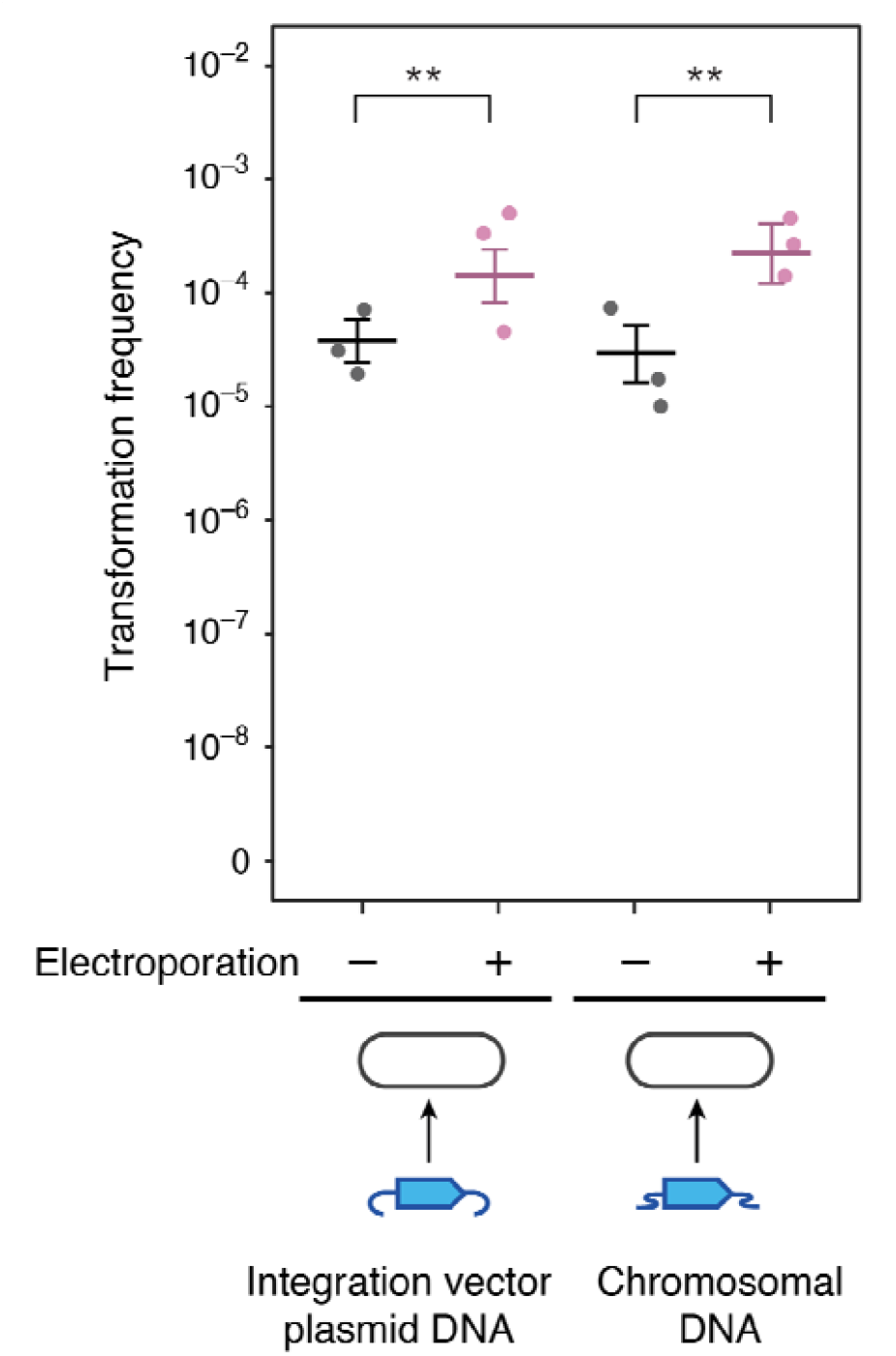
Effect of electroporation on *S. alvi* transformation. Competent cells were tested with and without electroporation to determine whether it led to a higher transformation frequency over the baseline level observed due to natural competence. Two types of DNA constructs were tested: integration vector plasmid DNA and chromosomal DNA. Statistical significance for comparisons: \*\**p* < 0.01; \**p* < 0.05; ns, nonsignificant (Poisson regression likelihood ratio tests).

**Supplementary Table 1 | Genetic changes in *S. alvi* after passaging through bees**

**Supplementary Table 2 | *S. alvi* competence gene homologs**

**Supplementary Table 3 | Bacterial strains used in this study**

